# Primary brain cell infection by *Toxoplasma gondii* reveals the extent and dynamics of parasite differentiation and its impact on neuron biology

**DOI:** 10.1101/2021.03.15.435467

**Authors:** Thomas Mouveaux, Emmanuel Roger, Alioune Gueye, Fanny Eysert, Ludovic Huot, Benjamin Grenier-Boley, Jean-Charles Lambert, Mathieu Gissot

**Affiliations:** Univ. Lille, CNRS, Inserm, CHU Lille, Institut Pasteur de Lille, U1019 - UMR 9017 - CIIL - Center for Infection and Immunity of Lille, F-59000 Lille, France; Univ. Lille, Inserm, Institut Pasteur de Lille, U1167, F-59000 Lille, France

## Abstract

*Toxoplasma gondii* is a eukaryotic parasite that forms latent cysts in the brain of immunocompetent individuals. The latent parasite infection of the immune-privileged central nervous system is linked to most complications. With no drug currently available to eliminate the latent cysts in the brain of infected hosts, the consequences of neurons’ long-term infection are unknown. It has long been known that *T. gondii* specifically differentiates into a latent form (bradyzoite) in neurons, but how the infected neuron responds to the infection remains to be elucidated. We have established a new *in vitro* model resulting in the production of mature bradyzoite cysts in brain cells. Using dual, host and parasite RNA-seq, we characterized the dynamics of differentiation of the parasite, revealing the involvement of key pathways in this process. Moreover, we identified how the infected brain cells responded to the parasite infection revealing the drastic changes that take place. We showed that neuronal-specific pathways are strongly affected, with synapse signaling being particularly affected, especially glutamatergic synapse signaling. The establishment of this new *in vitro* model allows investigating both the dynamics of parasite differentiation and the specific response of neurons to the long-term infection by this parasite.

## INTRODUCTION

*Toxoplasma gondii* is a unicellular eukaryotic pathogen. It belongs to the Apicomplexan phylum, which encompasses some of the deadliest pathogens of medical and veterinary importance, including *Plasmodium* (the cause of malaria), *Cryptosporidium* (responsible for cryptosporidiosis), and *Eimeria* (causative agent of coccidiosis)*. T. gondii* is an obligate intracellular parasite. Although toxoplasmosis is generally asymptomatic, it can lead to the development of focal central nervous system (CNS) infections in immunocompromised hosts. In addition, *Toxoplasma* is also a clinically important opportunistic pathogen that can cause birth defects in the offspring of newly infected mothers. The worldwide seroprevalence of *T. gondii* infection is estimated between 30 and 70% in humans, although it differs significantly depending on the geographical areas [1].

The life cycle of *T. gondii* is complex, with multiple differentiation steps that are critical to parasite survival in human and feline hosts [2]. Infection by oocysts containing sporozoites shed by cats or by bradyzoites contaminating ingested meat leads to differentiation into the rapidly growing tachyzoites that are responsible for the clinical manifestations in humans. The conversion of the tachyzoites into bradyzoites, responsible for the acute or the chronic phase of the disease, respectively, is made possible by the unique ability of the tachyzoite to spontaneously differentiate into the bradyzoite form in specific cell types such as muscle cells or neurons. These latent bradyzoites are thought to persist in the infected host for prolonged periods due to their ability to evade the immune system and to resist commonly used drug treatments. Bradyzoites have also the ability to reactivate into virulent tachyzoites and cause encephalitis, in particular in immunocompromised hosts [3]. Therefore, tachyzoite to bradyzoite interconversion is a critical step for the pathogenesis and survival of the parasite. *T. gondii* tachyzoite to bradyzoite stress-induced differentiation has been extensively studied *in vitro* using alkaline stress and other stimuli [4]. However, this process does not produce persisting cysts that express mature bradyzoite markers [5]. It merely reflects the complexity of the process observed *in vivo*. For example, much higher rates of spontaneous differentiation are observed in primary neurons [6]. However, infection of primary neurons was only performed for short periods (up to 4 days) [7–10]. Therefore, a global understanding of the kinetics and dynamics of differentiation is lacking due to widespread use of the imperfect, but easy to handle, stress-induced differentiation model.

*T. gondii* latent infection of the immune-privileged CNS is linked to most complications that can be fatal in the case of reactivation of bradyzoite cysts in immune-deficient hosts. These intracellular parasites migrate to the brain and cross the blood-brain barrier (BBB) by a *Trojan* horse mechanism [11] or by compromising the permeability of the BBB after infection and lysis of epithelial cells [12]. After reaching the CNS, the parasites can invade all nucleated cells, although infection is detected and persist in neurons *in vivo* [13]. Consistent with the ability of this parasite to infect and persist in neurons, *T. gondii* has been linked to behavioral changes in rodent models. The most prevalent study reported the ability of the parasite to specifically manipulate the behavior of rodents in relation to predator-prey interactions. In these studies, chronically infected mice were specifically impaired for their aversion to feline urine scent [14, 15]. Moreover, *T. gondii* infection has been directly implicated in modulating dopamine production [16], decreasing levels of norepinephrine and glutamate [17, 18], altering GABAergic signaling [19], thereby inducing an imbalance in neuronal activity [20], inducing neuron apoptosis [21] and altering synaptic protein composition [22]. Chronic toxoplasmosis is also correlated with the establishment of low-grade neuroinflammation characterized by the production of proinflammatory cytokine interferon-gamma (IFN-g). IFN-g is critical to control parasite replication [23] by inducing cell-autonomous immunity of immune resident brain cells notably astrocytes and microglia. Recently, *T. gondii-induced* neuroinflammation has also been linked to behavioral changes in rodents [24, 25] indicating that infection likely causes direct and indirect effects on neuronal functions. In humans, a growing number of studies have linked *T. gondii* to psychiatric diseases such as schizophrenia [26, 27], behavior alterations [28], and neurodegenerative diseases such as Parkinson and Alzheimer disease [29] although the causality is not direct and the effect of *T. gondii* infection on human behavior is likely subtle [30]. Indeed, chronic neuroinflammation may also cause neurological disorders either by producing neurodegeneration, neurotransmitter abnormalities and therefore altering the neuron functionality [31]. *T. gondii* infection may therefore have lifelong effects on the CNS of immunocompetent hosts.

Although global measurement of alteration at the whole brain level [32, 33] clearly indicates broad changes in neuron biological functions, the extent of the modifications of the individual neuron during long-term infection is not understood. Similarly, *in vivo* studies could not address the kinetics of the spontaneous differentiation of the parasite. To address this question, we reasoned that an *in vitro* culture of neurons would require the support of other cells such as astrocytes, which provide metabolic support for neurons and promote the function of synapses [34]. Therefore, we infected a complex primary brain cell culture with *T. gondii* tachyzoites to study the spontaneous differentiation dynamics and the host cell response to infection during differentiation and once the cysts are established. We show here that spontaneous differentiation occurs using this *in vitro* system and can be maintained for at least 14 days. Using RNA-seq, we characterized the dynamic changes in both parasite and host cell gene expression. We investigated the kinetics of parasite differentiation and the alteration of the brain cell gene expression after infection. We showed that this model produced infective bradyzoite cysts after 2 weeks of culture mirroring *in vivo* models. Thus, the *in vitro* model we established offers a unique opportunity to dissect the molecular mechanisms of parasite differentiation and the consequences of *T. gondii* infection on neuron biology.

## MATERIAL AND METHODS

### Parasite strains and culture

*Toxoplasma gondii* tachyzoites of the 76K strain were propagated *in vitro* in human foreskin fibroblasts (HFF) using Dulbeccos’s modified Eagles medium supplemented with 10% fetal calf serum (FCS), 2mM glutamine, and 1% penicillin-streptomycin. Tachyzoites were grown in ventilated tissue culture flasks at 37°C and 5% CO2. Prior to infection, intracellular parasites were purified by sequential syringe passage with 17-gauge and 26-gauge needles and filtration through a 3-μm polycarbonate membrane filter (Whatman)

### Brain cell culture

Animal housing and experimentation were carried out per the French Council in Animal Care guidelines for the care and use of animals and following the protocols approved by the Institut Pasteur de Lille’s ethical committee. Primary neuronal cultures were obtained from the hippocampus of postnatal (P0) rats as described previously [35]. Briefly, after the dissection of the brains, hippocampi were washed three times in HBSS (HBSS, 1-M HEPES, penicillin/streptomycin, and 100-mM sodium pyruvate, Gibco) and were dissociated via trypsin digestion (2.5%, 37°C, Gibco) for 7min. Next, hippocampi were incubated with DNase (5mg/mL, Sigma) for 1min and washed again in MEM medium supplemented with 10% SVF, 1% Glutamax, 0.8% MEM vitamins, 0.5% penicillin/streptomycin, and 0.45% d-glucose (Sigma). With a pipette, hippocampi were mechanically dissociated and resuspended in Neurobasal A, a medium supplemented with GlutaMAX and B27 neural supplement with antioxidants (Gibco). Cells were resuspended in culture medium, counted, and plated at a density of 100 000 cells/cm2 24-well plates. Plates were pre-coated with 0.1 □mg/ml poly-l-lysine in 0.1 □M borate buffer (0.31% boric acid, 0.475% sodium tetraborate, pH = 8.5; Sigma) overnight at 37°C and rinsed thoroughly with water. In total, 200,000 brain cells were seeded per well in 24-well plates. Brain cells were maintained at 37°C in a humidified 5% CO2 incubator. Brain cells were grown for 14 days before infection.

### Brain cell culture infection

Tachyzoites of the 76K strain were collected from an infected HFF T25 flask and purified by sequential syringe passage with 17-gauge and 26-gauge needles and filtration through a 3-μm polycarbonate membrane filter (Whatman). Brain cells that were grown and matured in Neurobasal A medium for 14 days were infected by the parasite. For that, the correct amount of tachyzoites was resuspended in 50 μL of Neurobasal A medium and then added onto the brain cell culture. Approximately 2.10^5^ brain cells were present in a well of a 24-well plate. Each well was infected by 3.10^4^ tachyzoites to a multiplicity of infection of 1 parasite for around 7 cells. The infected culture was maintained at 37°C in a humidified 5% CO2 incubator for the duration of the experiment without adding media to avoid disturbing the brain cell culture. A typical experiment yielded 2.5×10^4^ cysts per well of a 24-well plate (around 12.5 % of the 2.10^5^ brain cells).

### Mouse infection

Animal housing and experimentation were carried out in accordance with the French Council in Animal Care guidelines for the care and use of animals and following the protocols approved by the Institut Pasteur de Lille’s ethical committee (number #11082-2017072816548341 v2). Brain cells were infected as described above for a duration of 7 or 14 days. Infected and uninfected cells from a single well of a 24-well plate were scraped from the plates and resuspended in 400 μL of sterile PBS. Mice were gavaged with 200 μL of the solution containing the resuspended cells. The content of a single well of a 24-well plate was used to gavage two mice. Uninfected brain cell culture samples were collected at the same time as the 14 day-infected cells. Four weeks after gavage, brains were collected and homogenized individually. Cysts were counted after *Dolichol biflorus* lectin labeling of the cyst wall for 30 min at room temperature to a dilution of 1:400 in PBS. One fifth of the brain of each mouse was scored for the presence of lectin-positive cysts.

### RNA sample collection and library preparation

RNA samples were collected after infecting the primary brain cell cultures by the 76K strain for 24, 48, 96 hours, 7, and 14 days. Uninfected brain cell culture samples were collected at the same time as the 24 hours infected cells time-point. Infected and uninfected cells were washed with 1ml of PBS (2 times) and lysed by a direct load of Trizol in the plate. RNA was extracted as per manufacturer instruction and genomic DNA was removed using the RNase-free DNase I Amplification Grade Kit (Sigma). All RNA samples were assessed for quality using an Agilent 2100 Bioanalyzer. RNA samples with an integrity score greater than or equal to 8 were included in the RNA library preparation. Triplicates (biological replicates) were produced for each condition. The TruSeq Stranded mRNA Sample Preparation kit (Illumina) was used to prepare the RNA libraries according to the manufacturer’s protocol. Library validation was carried out by using DNA high-sensitivity chips passed on an Agilent 2100 Bioanalyzer. Library quantification was carried out by quantitative PCR (12K QuantStudio).

### RNA-seq and analysis

Clusters were generated on a flow cell within a cBot using the Cluster Generation Kit (Illumina). Libraries were sequenced as 50 bp-reads on a HiSeq 2500 using the sequence by synthesis technique (Illumina). HiSeq control software and real-time analysis component were used for image analysis. Illumina’s conversion software (bcl2fastq 2.17) was used for demultiplexing. Datasets were aligned with HiSAT2 v2.1.0 [36] against the *T. gondii* ME49 genome from (ToxoDB-39) [37] and against the rat genome (*Rattus norvegicus* Rn6 (UCSC)). Expression for annotated genes was quantified using htseq-count and differential expression was measured by DESeq2. P-values for multiple testing were adjusted using the Benjamini-Hochberg method. Differentially expressed genes with adjusted p-values below 0.05 and log2 fold-changes above 2 were considered in this study. Gene ontology was performed using the PANTHER [38] (Version 15) Overrepresentation Test (Released 20190711) surveying GO Slim Biological pathways using the Fisher statistical test for significance. RNA-seq data that support the findings of this study have been deposited in the GEO database under the accession number GSE168465.

### Immunofluorescence analysis

Infected and uninfected brain cell cultures were fixated using 4% PFA for 30 minutes. The coverslips were incubated with primary antibodies and then secondary antibodies coupled to Alexa Fluor-488 or Alexa-Fluor-594. Primary antibodies used for IFAs include anti-TgEno2, anti-TgSAG1, anti-MAP2, and anti-GFAP and were used at the following dilutions 1:1000, 1:1000, 1:500, and 1:500, respectively. A lectin from *Dolichos biflorus* coupled to fluorescein was also used at 1:400 dilution to identify the parasitic vacuoles. Confocal imaging was performed with a ZEISS LSM880 Confocal Microscope. All images were processed using Carl Zeiss ZEN software. Quantification of immunofluorescence assays was carried out manually by counting the concerned signal by visual observation. The signal corresponding to at least 100 vacuoles was counted for each replicate.

### Western-Blot

Total protein extracts representing infected or uninfected cells were resuspended in 1X SDS buffer. The protein samples were then fractionated on a 10% SDS-polyacrylamide electrophoresis gel and then transferred onto a nitrocellulose membrane. The anti-VGLUT1 (cat # 48-2400, Thermo-Fischer) and anti-GAPDH antibodies were used at a 1:1000 dilution. Chemiluminescent detection of bands was carried out by using Super Signal West Femto Maximum Sensitivity Substrate.

## RESULTS

### Establishment of the in vitro infection model of primary brain cell culture

To produce the primary brain cell culture, we extracted brain cells from newborn rats and placed them in culture for 14 days before infection. By immunofluorescence and after quantification, we determined that neurons represented at least 30 % of the cells present in culture as identified by the MAP2 marker (Figure S1A). Astrocytes, as identified by the GFAP marker, represent more than 50 % of the total cells while glial cells and oligodendrocytes represented around 20 % of all the cells (Figure S1A). This percentage did not vary over time (Figure S1A) or after infection (Figure S1B). Infection occurred and persisted in neurons and astrocytes and was maintained over time with a similar percentage of cells being infected until the 14 day time point (Figure 1A). To characterize the *T. gondii* spontaneous differentiation dynamics in this *in vitro* model, we followed the expression of tachyzoite (TgSAG1) and bradyzoite (Cyst wall labeled by *Dolichos bifluorus* lectin and p21, a late bradyzoite marker [39]) markers over time. Spontaneous differentiation occurred within a short time frame in the brain cells with the appearance of parasites expressing a marker of the cyst wall (labeled by the *D. bifluorus* lectin) 24h after infection representing more than 90 % of the parasite population after 96h (Figure 1B). Parasites expressing the tachyzoite marker TgSAG1 followed a reverse trend (Figure 1C). We noted the appearance of the late bradyzoite marker (p21) in cysts 96h after infection and more than 70 % of the cyst population was positive for this marker after 7 days (Figure 1D). Interestingly, we observed transitioning parasites until 48h of infection (expressing both tachyzoite and bradyzoite markers TgSAG1 and *D. bifluorus* lectin; Figure 1E), while all the parasites expressing p21 were also positive for the *D. bifluorus* lectin (Figure S1C). Imaging of parasites at 7 days after infection demonstrates that the parasites converted to bradyzoites and established latency in both astrocytes and neurons in this *in vitro* model (Figure 1F).

**Figure 1:**
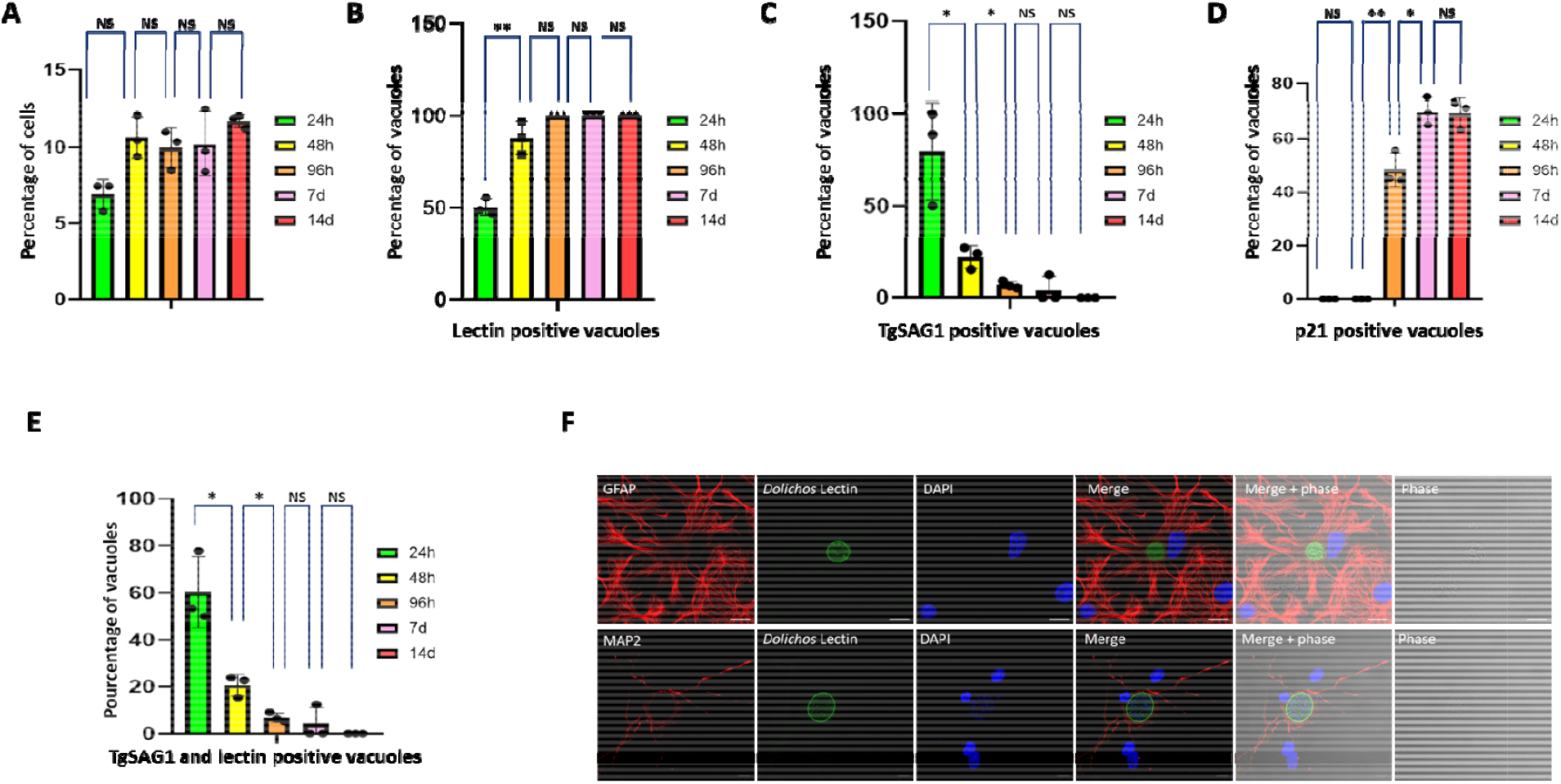
Critical aspects of the primary brain cell culture and its infection by *T. gondii*. **A**: Graphical representation of the number of infected cells in the brain primary cell culture. Bar graph representing the percentage of infected cells over time after 24h (green), 48h (yellow), 96h (orange), 7 days (pink) and 14 days (red) of infection. A Student’s t-test was performed; two-tailed p-value; NS: p>0,05; mean ± s.d. (n=3 independent experiments). **B**: Graphical representation of the number of *D. bifluorus* lectin positive vacuoles. Bar graph representing the percentage of infected cells over time after 24h (green), 48h (yellow), 96h (orange), 7 days (pink) and 14 days (red) of infection. A Student’s t-test was performed; two-tailed p-value; **: p<0,01; NS: p>0,05; mean ± s.d. (n=3 independent experiments). **C**: Graphical representation of the number of vacuoles expressing the tachyzoite marker TgSAG1. Bar graph representing the percentage of TgSAG1 positive parasite vacuoles over time after 24h (green), 48h (yellow), 96h (orange), 7 days (pink), and 14 days (red) of infection. A Student’s t-test was performed; two-tailed p-value; *: p<0,05; NS: p>0,05; mean ± s.d. (n=3 independent experiments). **D**: Graphical representation of the number of vacuoles expressing the late bradyzoite marker p21. Bar graph representing the percentage of p21 positive parasite vacuoles over time after 24h (green), 48h (yellow), 96h (orange), 7 days (pink) and 14 days (red) of infection. A Student’s t-test was performed; two-tailed p-value; *: p<0,05; **: p<0,01; NS: p>0,05; mean ± s.d. (n=3 independent experiments). **E**: Graphical representation of the number of vacuoles expressing both the tachyzoite marker TgSAG1 and presenting a lectin labeling. Bar graph representing the percentage of parasite vacuoles double-positive for TgSAG1and *D. bifluorus* lectin labeling over time after 24h (green), 48h (yellow), 96h (orange), 7 days (pink), and 14 days (red) of infection. A Student’s t-test was performed; two-tailed p-value; *: p<0,05; NS: p>0,05; mean ± s.d. (n=3 independent experiments). **F:** Immunofluorescence labeling of bradyzoite cysts in astrocytes and neurons 7 days post-infection. Confocal imaging demonstrating the presence of bradyzoite cysts (green, labeled with the *D. bifluorus* lectin) in astrocytes (upper panel, red, labeled with GFAP) or neurons (lower panel, red, labeled with MAP2). Anti-GFAP and anti-MAP2 were used as astrocyte and neuron markers, respectively. The scale bar (10 μm) is indicated on the lower right side of each confocal image.

### Dual RNA-seq on the parasite and host cell during the spontaneous parasite differentiation

To assess the transcriptome changes during the parasite spontaneous differentiation and the host response to infection, we collected triplicate RNA samples of infected primary CNS cell culture at 1d, 2d, 4d, 7d, and 14d post-infection (Figure 2A). We analyzed transcriptomic profiles of both the parasite and host cells (Figure S2A and S2B). Sequencing reads were assigned to the rat or the parasite genome (Table 1). For each time point, the infected host transcriptome was compared to a non-infected host cell culture. Reads assigned to the parasite genome were compared to purified tachyzoites derived sequencing reads. We used a p-value cut-off of 0,05 and a minimum 2-fold change to identify differentially expressed genes (DEG) using the DESEQ2 program (Table 2, Table S1 and S2). We performed a principal component analysis (PCA) to identify how each condition was clustering (Figure 2B and 2C). On the parasite side, the PCA analysis revealed that expression was similar between the time points 1d and 2d, while 4d appeared to represent the transition from the tachyzoite to the bradyzoite-specific expression observed at day 7d and 14d (Figure 2B). On the host side, PCA showed that the response to infection was different for the 1d and 2d time points compared to 7d and 14d (Figure 2C).

**Figure 2:**
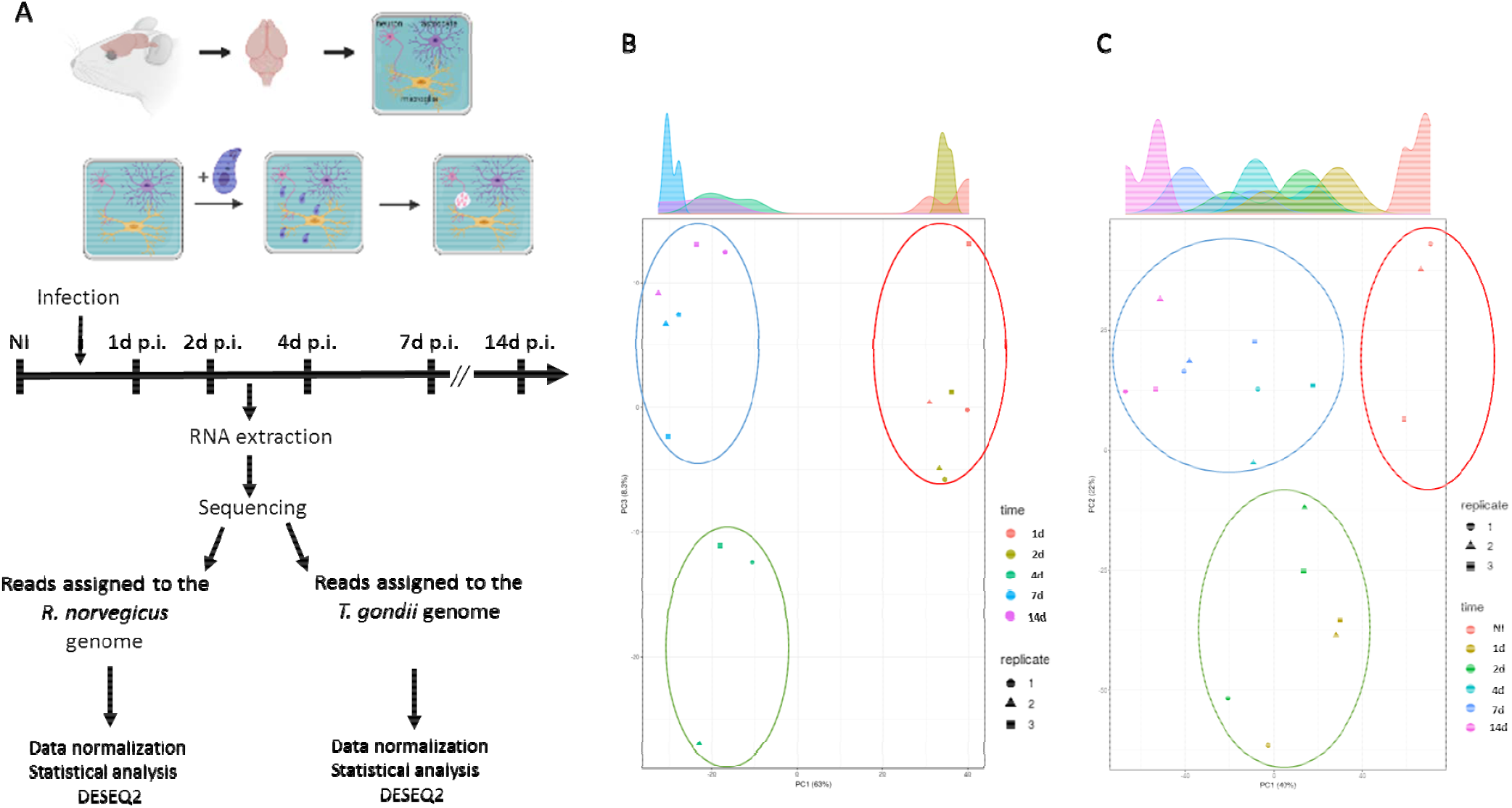
Dual RNA-seq on the uninfected and *T. gondii* infected primary brain cell culture. **A:** Schematic of the experiment representing the main steps of the primary brain cell culture and the time points when RNA was extracted. Libraries were created and processed through high throughput sequencing. Reads were assigned to either the *R. norvegicus* or *T. gondii* genome and DEGs were assigned using DESEQ2. **B:** Principal component analysis of the *T. gondii* triplicate results for each time point. Each replicate is represented by a square, a triangle, and a circle. Each time point was assigned a color: orange (1d), brown(2d), dark green (4d), dark blue (7d), and pink (14d). Based on this analysis, three main groupings were found and represented by a circle: red circle (1d and 2d), green circle (4d), and blue circle (7d and 14d) suggesting sharp transition during differentiation. **C:** Principal component analysis of the *R. norvegicus* triplicate results for each time point. Each replicate is represented by a square, a triangle, and a circle. Each time point was assigned a color: orange (non-infected, NI), brown (1d), green (2d), light blue (4d), dark blue (7d), and pink (14d). Based on this analysis, three main groupings were found and represented by a circle: red circle (non-infected, NI), green circle (1d and 2d), and blue circle (4d, 7d, and 14d).

**Table 1:** Number of reads assigned to the *R. Norvegicus* or *T. gondii* genes.

| Time point (days) | infection | Number of reads assigned to the rat genes | % Rat | Number of reads assigned to the *T.gondii* genes | % *T. gondii* | Number total of reads |
| --- | --- | --- | --- | --- | --- | --- |
| 1 | infected | 25252078 | 90.73 | 2579227 | 9.27 | 27831305 |
| 1 | infected | 22971803 | 94.27 | 1397078 | 5.73 | 24368881 |
| 1 | infected | 25176593 | 96.21 | 990451 | 3.79 | 26167044 |
| 2 | infected | 17660544 | 64.35 | 9783780 | 35.65 | 27444324 |
| 2 | infected | 19808387 | 79.82 | 5007576 | 20.18 | 24815963 |
| 2 | infected | 20621012 | 80.37 | 5036121 | 19.63 | 25657133 |
| 4 | infected | 18187041 | 64.73 | 10456789 | 35.27 | 28643830 |
| 4 | infected | 17572568 | 74.21 | 6106382 | 25.79 | 23678950 |
| 4 | infected | 22141916 | 80.89 | 5232408 | 19.11 | 27374324 |
| 7 | infected | 17699964 | 64.02 | 9949129 | 35.98 | 27649093 |
| 7 | infected | 16120663 | 68.96 | 7257703 | 31.04 | 23378366 |
| 7 | infected | 21899771 | 85.98 | 3569992 | 14.02 | 25469763 |
| 14 | infected | 23960501 | 84.48 | 4401325 | 15.52 | 28361826 |
| 14 | infected | 23324543 | 90.36 | 2488854 | 9.64 | 25813397 |
| 14 | infected | 23134477 | 91.05 | 2273878 | 8.95 | 25408355 |
|  | non infected | 924580 | 99.99 | 3214 | 0.01 | 927794 |
|  | non infected | 337050 | 99.99 | 3148 | 0.01 | 340198 |
|  | non infected | 160965 | 99.99 | 1398 | 0.01 | 162363 |
|  | tachyzoites | 1452 | 0,01 | 8825308 | 99.99 | 8826760 |
|  | tachyzoites | 1215 | 0,01 | 8085704 | 99.99 | 8086919 |
|  | tachyzoites | 966 | 0,01 | 8687601 | 99.99 | 8688567 |

**Table 2:** Number of identified DEGs for *R. norvegicus* and *T. gondii*.

| Comparison | Identified genes | DESeq2 DEG | Up | Down | Total cut-off >2 |
| --- | --- | --- | --- | --- | --- |
| ***R. norvegicus*** |  |  |  |  |  |
| 1d vs NI | 12 936 | 4642 | 1066 | 1523 | 2589 |
| 2d vs NI | 12 978 | 3362 | 1050 | 1184 | 2234 |
| 4d vs NI | 12 960 | 3114 | 837 | 784 | 1621 |
| 7d vs NI | 12 985 | 3995 | 1150 | 1232 | 2382 |
| 14d vs NI | 13 055 | 4907 | 1434 | 1690 | 3124 |
| ***T. gondii*** |  |  |  |  |  |
| 1d vs Tachyzoites | 7 212 | 2203 | 610 | 305 | 915 |
| 2d vs Tachyzoites | 7 381 | 3241 | 842 | 416 | 1258 |
| 4d vs Tachyzoites | 7 745 | 4634 | 1220 | 1005 | 2225 |
| 7d vs Tachyzoites | 7 768 | 5002 | 1843 | 1224 | 3067 |
| 14d vs Tachyzoites | 7 697 | 4459 | 1749 | 1035 | 2784 |

### Spontaneous parasite differentiation transition is reflected by specific expression patterns

We compared the parasite expression profiles obtained for each time point of the brain cell infected culture (Figure 3A). Differential expression mirrors the timing of spontaneous differentiation. Indeed, most of the changes are initiated at 1d and 2d p.i. and are maintained during later time points (Figure 3A, 636 DEG). At these time points, parasites are still transitioning (Figure 3A, Table 2, and Table S1). A turning point is observed at 4d post-infection when the late bradyzoite markers are detected in the *in vitro* culture (Figure 1), and parasites further differentiate to mature bradyzoites at days 7 and 14 (Figure 3A, 1200 DEG common to 4d, 7d, and 14d). Little changes are identified in the parasite transcriptome between day 7 and day 14 (Figure 3A, Table S1). The list of common DEGs between each time point encompasses the main bradyzoite markers such as BAG1, ENO1, LDH2, and BRP1 (Table 3). In contrast, tachyzoite markers (LDH1, ENO2, and SAG1) were repressed with a different dynamic (Table 3). While SAG1 is already repressed 1d after infection, ENO2 and LDH1 were significantly repressed only after 4d of infection (Table 3). We performed pathway-enrichment analyses based on the 1200 common DEGs for the 4d, 7d, and 14d time points, and found that classical pathways known to be repressed such as translation are overrepresented (Figure S3). Similarly, the GO-enriched pathways based on the upregulated genes are in line with the carbohydrate metabolism switch known to happen during differentiation (Figure S3) [40].

**Figure 3:**
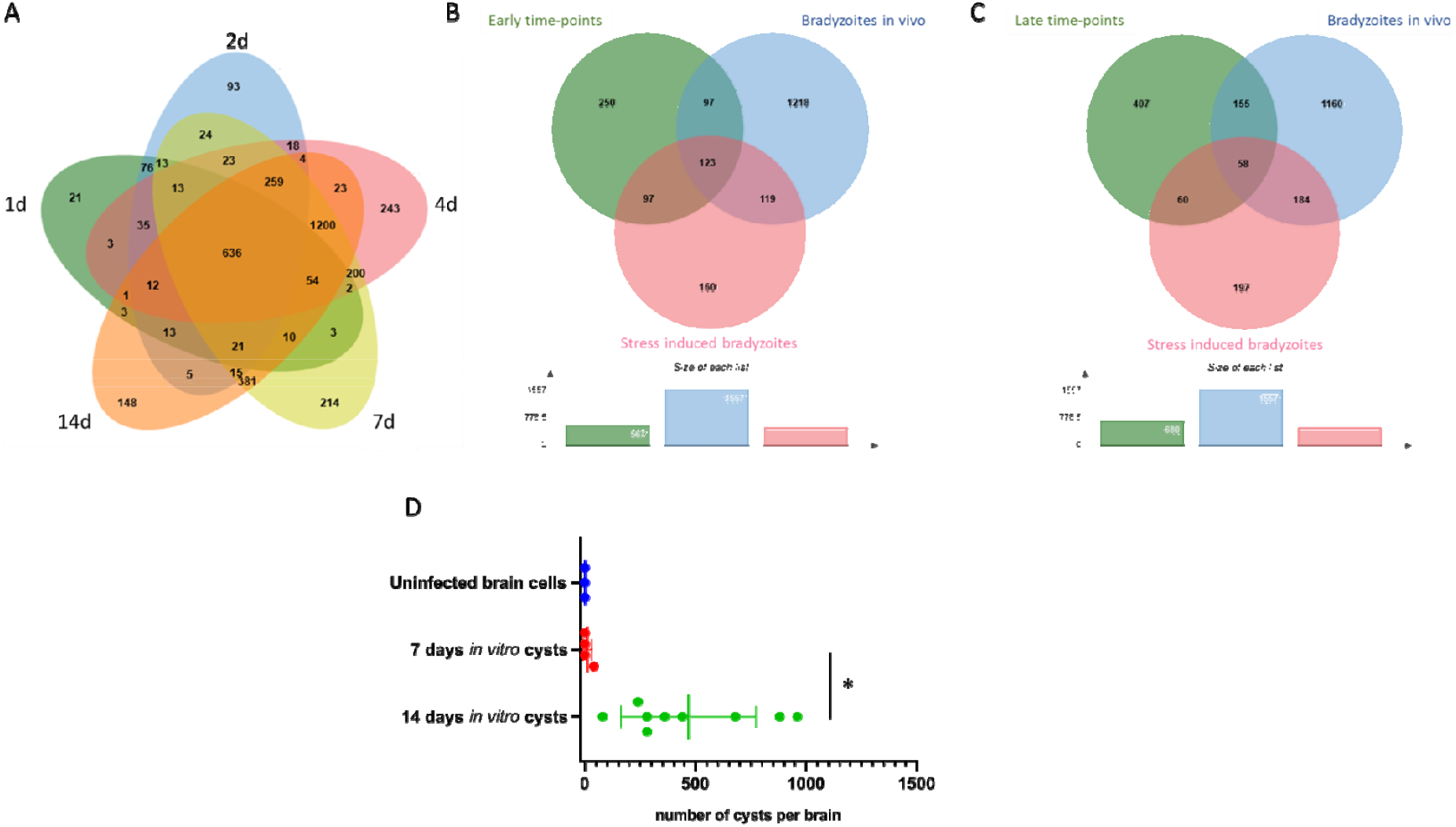
Bradyzoites produced in the infected primary brain cell culture are comparable to *in vitro* and *in vivo* produced bradyzoites. **A**: Venn diagram of the identified DEGs when comparing tachyzoite to parasite expressed genes at each time point of the brain cell culture. DEGs for the 1d time point are grouped in a green circle. DEGs for the 2d time point are grouped in a blue circle. DEGs for the 4d time point are grouped in a red circle. DEGs for the 7d time point are grouped in a yellow circle. DEGs for the 14d time point are grouped in an orange circle. Several unique or shared DEGs are indicated. **B**: Venn diagram of the identified upregulated DEGs common for the 1d and 2d time-points (green circle), the stress-induced upregulated DEGs (red circle) and the *in vivo* derived bradyzoites upregulated DEGs (blue circle). Some unique or shared DEGs are indicated. At the bottom, the size of each list of DEGs is indicated. **C**: Venn diagram of the identified upregulated DEGs common for the 4d, 7d, and 14d time-points (green circle), the stress-induced upregulated DEGs (red circle), and the *in vivo* derived bradyzoites upregulated DEGs (blue circle). Some unique or shared DEGs are indicated. At the bottom, the size of each list of DEGs is indicated. **D:** Bradyzoites cysts produced *in vitro* using the primary brain cell culture can transmit the infection after oral gavage. Mice were gavaged by uninfected (blue), 7 days (red) and 14 days (green) infected brain cells. After 6 weeks, mouse brains were collected and the number of cysts per brain was measured. A Student’s t-test was performed; two-tailed p-value; *: p<0,05;; mean ± s.d.

**Table 3:** Gene expression for tachyzoite and bradyzoite markers. Log_2_ fold change (FC) comparing the expression *T. gondii* transcripts at each time point of the infected brain cell culture to that of purified tachyzoites. Color gradient depends on the value of FC. Downregulated values are represented by shades of green. Upregulated values are represented in shades of red. For each transcript, the gene identification number (gene ID) and the corresponding annotation is also presented.

| Gene ID | Annotation | 1d | 2d | 4d | 7d | 14d |
| --- | --- | --- | --- | --- | --- | --- |
| TGME49_268860 | ENO1 | 5,48 | 7,34 | 10,00 | 10,50 | 10,17 |
| TGME49_291040 | LDH2 | 7,49 | 8,71 | 9,87 | 10,55 | 10,26 |
| TGME49_259020 | BAG1 | 6,65 | 8,76 | 9,64 | 10,29 | 10,28 |
| TGME49_314250 | BRP1 | 3,71 | 4,84 | 3,47 | 4,05 | 4,33 |
| TGME49_233460 | SAG1 | -1,91 | -2,37 | -4,65 | -7,29 | -6,48 |
| TGME49_268850 | ENO2 | -0,46 | -0,39 | -1,72 | -3,53 | -3,15 |
| TGME49_232350 | LDH1 | -0,53 | -0,65 | -1,54 | -2,04 | -1,78 |

### Parasites established in brain cell culture may represent bradyzoites

Expression profiles during stress-induced differentiation were already characterized in numerous studies [41–43]. We compared the expression profiles of up and down-regulated genes after alkaline stress-induced differentiation with the brain cell infected culture RNA-seq results. To account for experimental design and strain differences, we gathered a list of DEGs after alkaline stress-induced differentiation that was common to these three experiments [41–43]. We also compared our dataset to the DEGs that were identified after RNA-seq on *in vivo* derived bradyzoites [40]. Since there is a clear phenotypic switch between the early time-points (1d and 2d) of the infected brain cell culture and the late time-points (4d, 7d, and 14d), we extracted the DEGs that were common to either early time-points (1d and 2d) or late time-points (4d, 7d, and 14d). This comparison was carried out for upregulated DEGs (Figure 3B and 3C) and down-regulated DEGs (Figure S4A and S4B). At early time points, the number of shared up-regulated DEGs is equivalent between our dataset and the alkaline stress-induced differentiation or *in vivo* derived bradyzoites (Figure 3B). In contrast, at the late time points, the brain cell infected culture DEGs are closer to the *in vivo* derived bradyzoites DEGs than the alkaline stress-induced differentiation DEGs (Figure 3C). Similar results were obtained for the down-regulated genes (Figure S4A and S4B). This indicates that these late time-point brain cell produced bradyzoites may better represent the slow maturation of bradyzoites that is observed *in vivo*. However, brain cell-derived, *in vivo*-derived and stress-induced bradyzoites appear to be three distinct populations with regards to DEGs. Overall, brain cell-derived bradyzoites do not match the *in vivo* bradyzoite profile better than stressed induced-derived bradyzoites.

We investigated if the bradyzoite cysts produced *in vitro* using brain cells could be able to infect mice after oral gavage. In this experiment, the cysts have to go through the digestive system and release the bradyzoites in the gut of the mouse to proceed to the infection of intestine cells. The parasites will then turn into tachyzoites and eventually produce cysts in the brains. We used the cysts formed *in vitro* after 7 or 14 days of differentiation and uninfected brain cells to gavage mice. Six weeks after gavage, we collected the brains of the infected mice and probed for the presence of cysts. All the mice that were gavaged using 14 days *in vitro* cysts were successfully infected and presented cysts in their brain, while only one mouse presented cysts when using 7 days *in vitro* cysts (Figure 3D) indicating that 14 day cysts may have gone through more maturation steps. No cysts were found in the mice infected by brain cells alone (Figure 3D).

### Expression patterns during parasite differentiation suggest an overhaul of invasion and host-cell remodeling activities in the bradyzoite

Tachyzoites have a distinctive ability to modulate the expression of host cells by injecting parasite proteins to hijack the host’s regulatory pathways [44]. Very limited information is available about the expression of exported proteins from bradyzoites [45, 46] and their abilities to manipulate the host cells. We examined the expression of effector proteins that are known to be exported to the host cell cytosol and nucleus [44]. In our dataset, we found that most of the known effectors were downregulated during differentiation indicating that their expression is no longer needed for bradyzoite development (Table 4). Notably, TgIST was the only effector that presented a similar expression level in tachyzoites and bradyzoites and this was for all the time points examined (Table 4). As shown before for tachyzoite and bradyzoite markers (Table 3), day 4 represented a breaking point where the bradyzoite expression program replaces that of the tachyzoite. Exploring the expression of other potential effectors suggested that a complete transformation in the expression of these proteins is taking place during differentiation (Table S3). We also investigated the expression of proteins specialized in the invasion of host cells to verify if the bradyzoites also adapted their invasion machinery. Surprisingly, most of the proteins known to be important for tachyzoite invasion were downregulated (Table 5). Instead, a specialized subset of genes (RON2L1, RON2L2, sporoAMA1, AMA2, and AMA4 to a lesser extent) were over-expressed in bradyzoites especially at later time points. These proteins could potentially functionally replace in bradyzoites the tachyzoite specific AMA1 and RON2 proteins (Table 5). However, as shown previously in *in vivo* derived bradyzoites datasets [40], the reads’ coverage for sporoAMA1 is only partial in the late time points (14d) indicating that this gene likely produces truncated transcripts and proteins at that stage (Figure S5A).

**Table 4:** Gene expression for known tachyzoite effectors. Log_2_ fold change (FC) comparing the expression *T. gondii* transcripts at each time point of the infected brain cell culture to that of purified tachyzoites. Color gradient depends on the value of FC. Downregulated values are represented in shades of green. For each transcript, the gene identification number (gene ID) and the corresponding annotation is also presented.

| Gene ID | Annotation | 1d | 2d | 4d | 7d | 14d |
| --- | --- | --- | --- | --- | --- | --- |
| TGME49_308090 | ROP5 | -2,53 | -2,67 | -4,48 | -3,37 | -3,25 |
| TGME49_205250 | ROP18 | -0,97 | -0,97 | -2,85 | -2,28 | -2,31 |
| TGME49_258580 | ROP17 | -0,85 | -0,66 | -2,66 | -2,47 | -2,00 |
| TGME49_262730 | ROP16 | -0,34 | -0,47 | -3,28 | -3,66 | -3,58 |
| TGME49_275470 | GRA15 | -0,81 | -0,75 | -4,71 | -5,33 | -4,00 |
| TGME49_208830 | GRA16 | -2,00 | -1,61 | -4,00 | -4,75 | -4,52 |
| TGME49_230180 | GRA24 | -1,26 | -2,11 | -2,01 | -3,30 | -3,26 |
| TGME49_240060 | TgIST | 0,46 | 0,49 | 0,59 | 0,71 | 0,54 |

**Table 5:** Gene expression for transcripts encoding proteins known to be involved in invasion. Log_2_ fold change (FC) comparing the expression *T. gondii* transcripts at each time point of the infected brain cell culture to that of purified tachyzoites. Color gradient depends on the value of FC. Downregulated values are represented in shades of green. Upregulated values are represented in shades of red. Transcripts that were not detected are indicated by a double dash line (--). For each transcript, the gene identification number (gene ID) and the corresponding annotation is also presented.

| Gene ID | Annotation | 1d | 2d | 4d | 7d | 14d |
| --- | --- | --- | --- | --- | --- | --- |
| TGME49_255260 | AMA1 | -1,20 | -1,45 | -1,47 | -1,25 | -1,38 |
| TGME49_294330 | AMA4 | -1,35 | -0,58 | 0,31 | 0,05 | 0,52 |
| TGME49_300130 | AMA2 | 0,89 | 0,90 | 4,29 | 4,56 | 4,41 |
| TGME49_315730 | sporoAMA1 | -- | -- | 8,60 | 9,51 | 9,37 |
| TGME49_265120 | sporoRON2 - RON2L2 | 3,52 | 4,00 | 6,40 | 6,38 | 6,67 |
| TGME49_294400 | RON2L1 | 0,82 | 1,35 | 0,80 | 0,90 | 0,97 |
| TGME49_310010 | RON1 | -0,51 | -0,44 | -1,59 | -0,97 | -0,92 |
| TGME49_300100 | RON2 | -0,88 | -0,83 | -2,59 | -2,29 | -1,84 |
| TGME49_223920 | RON3 | -0,53 | -0,44 | -2,26 | -2,52 | -2,08 |
| TGME49_229010 | RON4 | -0,77 | -0,88 | -2,48 | -2,27 | -1,62 |
| TGME49_311470 | RON5 | -1,04 | -1,11 | -2,29 | -1,85 | -1,47 |
| TGME49_297960 | RON6 | -0,86 | -0,87 | -2,27 | -1,92 | -1,74 |
| TGME49_306060 | RON8 | -0,69 | -0,74 | -2,27 | -2,35 | -1,90 |
| TGME49_308810 | RON9 | -0,39 | -0,33 | -0,82 | -0,80 | -0,60 |
| TGME49_261750 | RON10 | -0,98 | -0,99 | -1,09 | 0,00 | -0,07 |

In contrast, the AMA2 gene seems to produce full-length transcripts that are preferentially expressed in the late time points of the brain cell infected culture (Figure S5B). In line with these profound changes, the expression pattern of ApiAP2 transcription factors that may be responsible for the establishment of the specific expression profile varied also during differentiation (Figure S6). ApiAP2 expression profiles grouped in different clusters (Figure S6A): a first bradyzoite cluster induced early during differentiation that contained AP2IX-9 [47], a second bradyzoite cluster with factors induced later during differentiation containing AP2XI-4 [48] and a tachyzoite specific cluster with AP2IX-5 [49] and AP2XI-5 and AP2X-5 [50]. Principal component analysis based on the ApiAP2 expression profiles mirrored the transition during differentiation (Figure S6B). ApiAP2 transcription factors that may control different processes during differentiation may be present in the bradyzoite cluster.

### Brain cell culture showed a differential response to tachyzoite and bradyzoite infection

On the host side, infection by *T. gondii* tachyzoites triggered a strong response of the host cells (Table 2 and Table S2). This response is mostly stable during the 14d of infection since a large number of DEGs are common between each time point (834 DEGs, Figure 4A and 4B). However, the early response at 1d (with 521 unique DEGs) and 2d (318 DEGs only present at day 1 and 2 p.i) may be specific to acute infection (Figure 4A). We also noted that the later time points (7d and 14d p.i) presented a unique differential expression pattern (531 DEGs specific from 14d and 433 only common to 7d and 14d). This indicates that a distinctive host response to tachyzoite infection (early time points) is induced when compared to the time when cysts are established (7 and 14 days p.i). We separated DEGs between upregulated (Figure S7A) and downregulated (Figure S7B) and we identified similar trends with a number of DEGs being shared between each time point and representing the common response to infection. We also noted that a subset of DEGs was upregulated or downregulated at the first time points while a specific response was also emerging for later time points.

**Figure 4:**
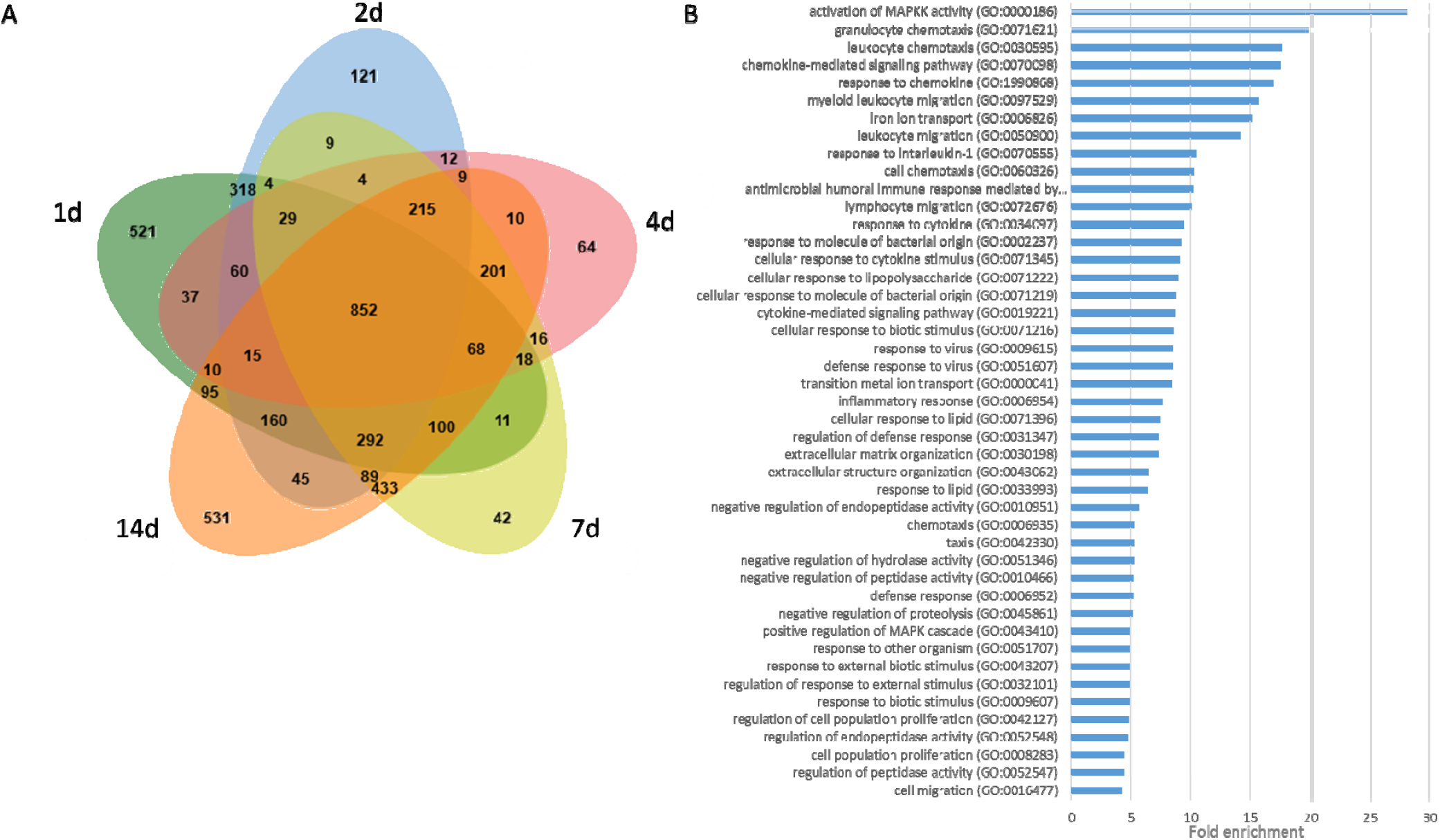
Analysis of identified *R.norvegicus* DEG in the infected primary brain cell culture when compared to uninfected samples. **A**: Venn diagram of the identified DEGs for each time point. DEGs for the 1d time point are grouped in a green circle. DEGs for the 2d time point are grouped in a blue circle. DEGs for the 4d time point are grouped in a red circle. DEGs for the 7d time point are grouped in a yellow circle. DEGs for the 14d time point are grouped in an orange circle. Several unique or shared DEGs are indicated. The total number of DEGs for each time point is indicated at the bottom of the figure. **B**: Enriched GO pathways for upregulated DEGs that are shared for all time-point infected brain cells. Pathways were selected with an FDR of 0,05 and a minimum enrichment of 4. The name of each GO pathway is indicated on the left part of the figure. Bars represent the enrichment fold.

### *Upregulation of immune-related pathways is a hallmark of* T. gondii *infected brain cell culture*

We performed a pathway enrichment analysis on the rat genes that are differentially expressed when comparing the brain cell uninfected cultures to the infected cultures at different time points (Figure 4B and 5A). First, we looked into up-regulated genes that were common for all time points and identified that the main response was an immune response to the infection that lasted during the 14 days of infection (Figure 4B). In particular, the response to chemokine (GO: 1990868) and the chemokine-mediated signaling pathway (GO:0070098) was overrepresented (Figure 4A). Similarly, upregulated DEGs belonging to the cellular response to cytokine stimulus (GO:0071345) and response to cytokine (GO:0034097) pathways were also overrepresented. Moreover, the response to interleukin-1 (GO:0070555) was also enriched in this dataset. This is in line with the neuroinflammation observed *in vivo* [51] and likely reflects the activation of astrocytes and glial cells present in the culture. This indicates that both microglia and astrocytes present in the brain cell culture responded strongly to the infection *in vitro*. Moreover, a specific response is observed in early time points (days 1 and 2), with a clear enrichment of genes involved in cell cycle and DNA replication arrest (GO:0045839 and GO:0051985) indicating that infection may induce an arrest of cell division of the brain cells such as glial cells (Figure S8A). At later time points, further activation of microglia may take place with the CD80 expression along with Galactin9 expression (Figure S8B).

**Figure 5:**
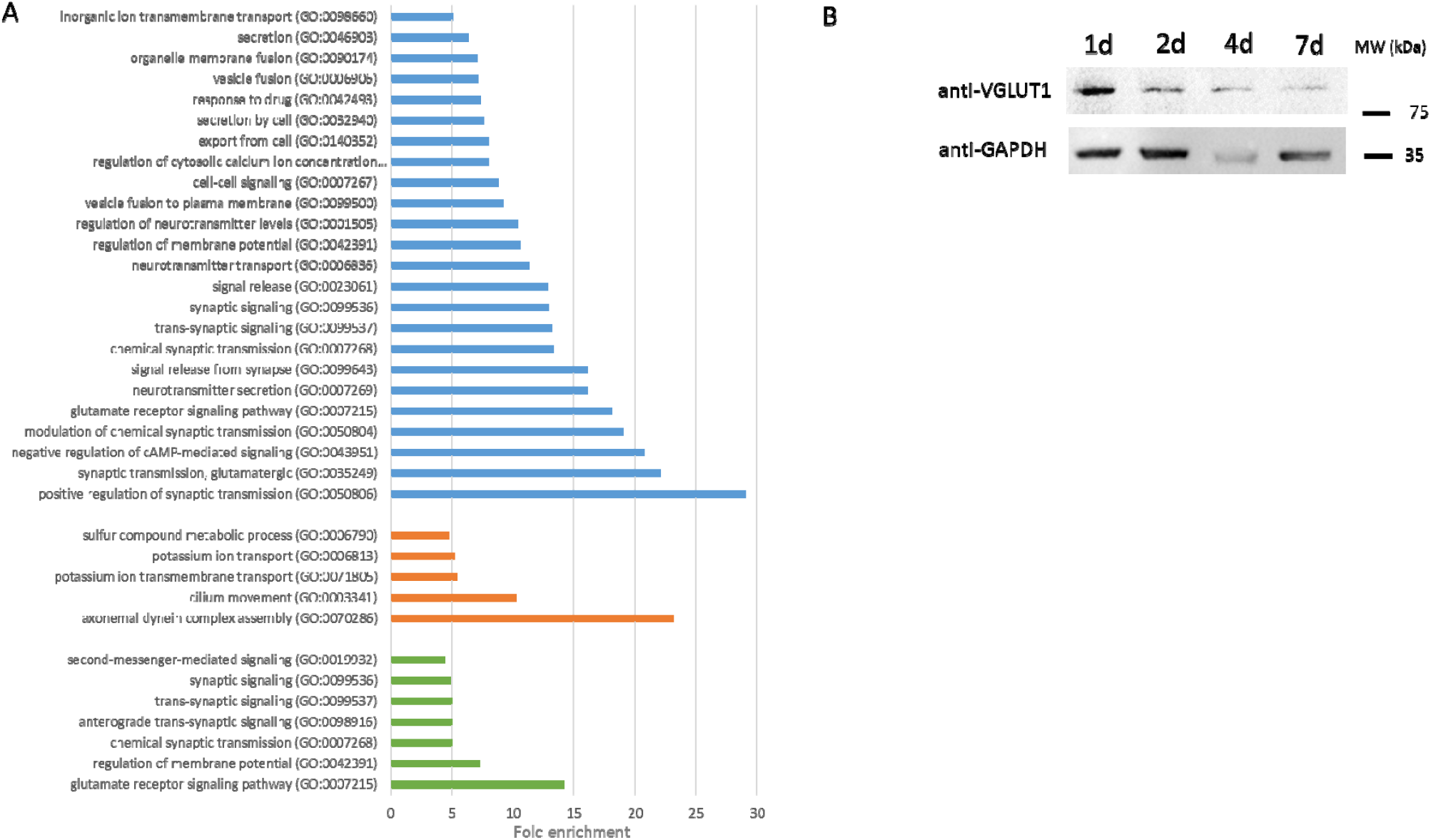
Gene ontology analysis of enriched downregulated pathways in brain cells. **A**: Enriched GO pathways for downregulated DEGs that are shared for all time-point infected brain cells (blue bars), shared for 1d and 2d time points (orange bars), and shared for 7d and 14d time points (green bars). Pathways were selected with an FDR of 0,05 and a minimum enrichment of 4. The name of each GO pathway is indicated on the left part of the figure. Bars represent the enrichment fold. **B:** Western blot showing the expression of Grm1 (VGLUT1) in neurons after 1, 2, 4, and 7 days of infection. GAPDH is used as a loading control.

### T. gondii *infection induces downregulation of key neuron functions and pathways*

Downregulated DEGs common to all time points were analyzed using gene ontology. The synapse function was impacted at all time points (Figure 5A). Notably, the most enriched pathways downregulated were linked to synapse plasticity and transmission (GO:0050804, GO:0007269, and GO:0007268). In particular, the glutamatergic synapse was affected with the down-regulation of metabotropic glutamate receptors (Grm1, 2, and 4) and Glutamate ionotropic receptor (Grik1, NMDA2C, and 2D) as previously described *in vivo* [18]. At later time points, the downregulation of a supplementary metabotropic glutamate receptor (Grm8) together with Homer 1 and 2 protein homologs that link the glutamate receptor to downstream signaling, indicated potential long-term impairment of the glutamate receptor signaling pathway (GO:0007215). We inspected the expression of the Grm1 protein during the infection of brain cells and confirmed the down-regulation of this protein illustrating the long-term effects of *T. gondii* infection on the glutamatergic synapse (Figure 5B and Figure S8C). Similarly, the glutamate decarboxylase isoforms (Gad1 and Gad2), responsible for GABA production in neurons, were downregulated since 1d recapitulating what was observed *in vivo* [19]. The synaptic signaling was also globally impacted with the downregulation of numerous membrane trafficking regulatory transcripts such as Synaptotagmin-1, Synapsin-2, or Otoferlin.

At early time points (1d and 2d), a specific response to infection consisted of the downregulation of axonemal dynein complex assembly (GO:0070286) pathway that suggested an arrest of axonemal assembly. At the same time points, the generation of the action potential and therefore excitability of neurons may be impacted by the downregulation of the potassium ion transmembrane transport (GO:0071805) pathway that may occur in neurons or astrocytes. Expression of both the regulatory membrane potential (GO:0042391) and chemical synaptic transmission (GO:0007268) pathways was also further decreased at late time points of infection suggesting a strong impact on neuron function.

## DISCUSSION

Tachyzoite to bradyzoite differentiation is a key aspect of *T. gondii* biology and pathogenesis. To date, it has been mainly tackled through the use of an *in vitro* model of stress-induced differentiation that merely reflected the process of spontaneous differentiation observed *in vivo*. Moreover, little is known on the consequences of the long-term infection of targeted host cells *in vivo* (mainly neuron and muscle cells). To better assess the spontaneous differentiation process and the host cell response to infection, we established a complex *in vitro* model where parasites are in contact with multiple cell types normally present in the brain. We reasoned that this complex environment will permit a sustainable long-term infection model. We were able to produce a viable environment promoting neuron survival for a minimum time of 28 days. Using this composite *in vitro* culture system, we successfully established and maintained the infection of neurons and astrocytes by the parasite that progressively express mature bradyzoite markers for at least 14 days. Primary neuronal infection by tachyzoites and bradyzoite differentiation was already experimented in different models for short time frames (up to 4 days) [7–10]. We were able to produce cysts in neurons that could be kept in culture for at least 14 days although longer time frames could be achieved (30 days, data not shown). Strikingly, the cysts produced using this new *in vitro* system have all the molecular features of mature cysts previously observed *in vivo*. They are also infective by oral gavage demonstrating that some of the cysts in the brain cell culture present an intact cyst wall and these *in vitro* produced bradyzoites can readily infect the mouse intestine. Surprisingly, bradyzoites were found in both neurons and astrocytes, a feature that is found in rat, mouse, and human primary brain cell culture [6, 9, 52] but not in mouse brains where bradyzoite survival is only sustained in neurons [13]. Immune cells, that are absent in the primary brain cell culture, may be crucial to eliminate the infected astrocytes *in vivo*.

We showed that parasite expression of bradyzoite markers appeared early in the differentiation process suggesting that the parasites are switching expression patterns at the beginning of the infection process. We observed parasites that were able to co-express markers of both tachyzoite and bradyzoite forms. This illustrates that differentiation is a dynamic process during which tachyzoites expressing bradyzoite markers can be observed until 4 days into the transition. RNA-seq also demonstrated that tachyzoite marker expression is only significantly repressed after 4 days. Such co-expression has also been observed during differentiation *in vivo* [53]. After 7 days, the expression profiles revealed by RNA-seq suggest that the parasites present in the brain cell culture have mainly switch to a bradyzoite specific expression program. We did not observe major differences in gene expression between 7 and 14 days of culture (Table S1). However, only the 14 day bradyzoites containing cysts were competent for mouse infection through gavage indicating that a maturation process, which is not reflected by transcriptional changes, is still undergoing after 7 days. This post-transcriptional maturation process may involve the modification of the cyst wall.

The parasites produced after 14 days of *in vitro* culture are therefore infectious by oral gavage. In this proof of principle experiment, we showed that using half of a single well of 24-well plate of brain cell derived bradyzoites is sufficient to produce cysts *in vivo* after oral gavage. However, more work is needed to establish how many brain-cell derived cysts are sufficient to infect a mouse. The *in vitro* culture model described here may be a way to reduce experimental mouse usage. The simplicity to produce the starting material (1 well of a 24 well-plate can be used to infect two mice) also offers the possibility to test the infectiousness by oral gavage of multiple parasite mutants. Interestingly, similar results were obtained using a human myotube-based *in vitro* culture model [54], indicating that *in vitro* production of infectious cysts is also possible in other cell types for which a tropism exists *in vivo*.

By examining the expression pattern of transitioning parasites, we observed that the expression of ApiAP2 transcription factors was differentially regulated. Two clusters that appeared early and late during differentiation were identified and may coordinate the dynamic expression profiles observed in the brain cell culture. Interestingly, the over-expression of BFD1, the master switch of differentiation [55], was only observed from 4 days onwards, although its expression might be regulated through a post-transcriptional mechanism. This indicates that multiple layers of regulation may be essential to produce mature bradyzoites.

We have also identified that the expression of the major tachyzoite effectors of host-cell manipulation was repressed during differentiation except for TgIST. This suggests that the bradyzoites express a new set of proteins to enable their persistence in neurons. It would be interesting to characterize the proteins that are specifically expressed during differentiation and that have the potential to be exported in the host cell. We also observed the same phenomenon for proteins known to be involved in invasion. Invasion proteins such as AMA1 and RON2, which are key to form a tight connection between the invading parasite and host cell membranes, may be replaced in the bradyzoites by AMA2 or AMA4 and RON2L1 or RON2L2. This modification may be necessary for the bradyzoites to invade specific host cells, such as enterocytes, to complete the life cycle. These new findings are critical for understanding the fundamental changes that occur after differentiation. It suggests that bradyzoites remodel their parasite-host interaction machinery to adapt to a narrower host cell range (intestine enterocyte, neurons, and muscle cells) compared to tachyzoites.

Neurons are strongly impacted by *T. gondii* infection. We found that both GABA and glutamate signaling were disrupted in the brain cell culture much like what has been observed *in vivo* in *T. gondii* infected mouse brains. The glutamate signaling is disrupted from the beginning of the infection with the downregulation of both metabotropic glutamate receptors and glutamate ionotropic receptors. The latter was shown to be repressed in mouse infected brains [56] and participate in a process proposed to contribute to the establishment of psychiatric disorders such as schizophrenia although the effect of *T. gondii* infection on human behavior is likely subtle [30]. Thus, this study extends the number of receptors that may be downregulated during infection and further emphasize the impact of infection and inflammation on glutamate signaling.

We also discovered that early on in infection, axonemal growth might be repressed. Development, as well as maintenance of correct cilia structure, is essential for the unique neuron sensory properties, suggesting that neurons may respond to infection by limiting their ability to transfer information. Repression of membrane trafficking regulatory mechanisms was also observed suggesting that the synapse function may be disrupted. This may be aggravated when the parasite established a long-term infection since both membrane potential and chemical synaptic transmission are further disturbed at later time points of the infection. Our data, expand and confirm the extent of neuronal function disruption during *T. gondii* infection.

*T. gondii* infection has been linked to a change in behavior in rodents [14, 15]. The strong disruption of glutamate and GABA signaling previously reported [19] is confirmed by our study and may provide a link between the behavior changes and the infection by *T. gondii*. Since we also observed a signature of a strong neuro-inflammation as was shown *in vivo*, it is difficult to define the contribution of the direct infection of neurons and the indirect effects of neuro-inflammation on the neuronal pathways. Recent data [24, 25] indicate the importance of neuro-inflammation in *T. gondii*-induced behavioral changes.

We have established that parasites spontaneously differentiate when infecting a primary brain cell culture. Differentiated parasites present the hallmarks of bradyzoites and persist in culture for prolonged periods. Therefore, this *in vitro* system provides a unique opportunity to dissect the dynamic features of parasite differentiation but also the direct effect of infection on neuron biology. It could also be of interest for the screening of novel molecules that may be able to eliminate the parasite cyst once it is established in the neurons.

## Acknowledgments

The authors wish to thank the BioImaging Center Lille for access to instruments and Dr. Marion and Asma S. Khelifa for critically reading the manuscript.

## Author contributions

TM: Data collection, Data analysis, and interpretation; ER: Critical revision of the article, Drafting the manuscript; AG: Data collection; FE: Data collection; LH: Data collection; BGB: Data analysis and interpretation; JCL: Conception or design of the work, Drafting the manuscript; MG: Conception or design of the work, Drafting the manuscript, Data analysis and interpretation.

