## Supplementary material for "Primary brain cell infection by *Toxoplasma gondii* reveals the extent and dynamics of parasite differentiation and its impact on neuron biology": Supp figures and tables legends

**Figure S1: Dynamics of the *T. gondii* infected and uninfected primary brain cell culture.**

**Figure S1A:** Bar graph representing the percentage of neurons (blue bars) and astrocytes (red bars) over time after 24h, 48h, 96h, 7 days and 14 days.

**Figure S1B:** Bar graph representing the percentage of neurons (blue bars) and astrocytes (red bars) over time after 24h, 48h, 96h, 7 days and 14 days after infection.

**Figure S1C:** Graphical representation of the number of vacuoles expressing both the tachyzoite marker TgSAG1 and presenting a lectin labelling. Bar graph representing the percentage of parasite vacuoles double positive for p21 and *D. bifluorus* lectin labelling over time after 24h (green), 48h (yellow), 96h (orange), 7 days (pink) and 14 days (red) of infection. A Student’s t-test was performed; two-tailed p-value; *: p<0,05; **: p<0,01; NS: p>0,05; mean ± s.d. (n=3 independent experiments).

**Figure S2: Scatter plot representation of raw reads counts.**

Figure S2A: Scatter plot representation of raw reads counts for *T. gondii*. Each sample is compared to all the other samples.

Figure S2B: Scatter plot representation of raw reads counts for *R. norvegicus*. Each sample is compared to all the other samples.

**Figure S3: Gene ontology analysis for the 1200 common DEG for the 4d, 7d and 14d time points**. Each enriched pathway is indicated at the bottom. The y axis represents the fold enrichment for each enriched GO pathway. The size of the circle represents the -log_10_ of the p-value. Downregulated GO pathways are indicated in blue. Upregulated GO pathways are indicated in red.

**Figure S4: Comparison of the downregulated DEGs in brain cells, after stress induced differentiation and in *in vivo* derived bradyzoites.**

**Figure S4A**: Venn diagram of the identified downregulated DEGs common for the 1d and 2d time-points (green circle), the stress-induced downregulated DEGs (red circle) and the *in vivo* derived bradyzoites downregulated DEGs (blue circle). Number of unique or shared DEGs are indicated. At the bottom, the size of each list of DEGs is indicated.

**Figure S4B**: Venn diagram of the identified downregulated DEGs common for the 4d, 7d and 14d time-points (green circle), the stress-induced downregulated DEGs (red circle) and the *in vivo* derived bradyzoites downregulated DEGs (blue circle). Number of unique or shared DEGs are indicated. At the bottom, the size of each list of DEGs is indicated.

**Figure S5: Analysis of the read coverage for SporoAMA1 and AMA2 in different samples**

Figure S5A: Read coverage for SporoAMA1. The read coverage for the triplicate samples of tachyzoites, 14d brain cell culture and from *in vivo* bradyzoite [40] samples is represented. The gene number is indicated at the top of the figure.

Figure S5A: Read coverage for AMA2. The read coverage for the triplicate samples of tachyzoites, 14d brain cell culture and from *in vivo* bradyzoite [40] samples is represented. The gene number is indicated at the top of the figure.

**Figure S6: Analysis of ApiAP2 transcription factors during differentiation.**

Figure S6A: Clustering of annotated ApiAP2 transcription factors based on their expression during the infection of brain cells. Two bradyzoite clusters were identified (in blue). One tachyzoite cluster is identified (in orange).

Figure S6B: Principal component analysis based on the expression of ApiAP2 transcription factors.

**Figure S7:** **Analysis of identified *R. norvegicus* upregulated and downregulated DEGs in the infected primary brain cell culture when compare to uninfected samples.**

Figure S7A: Venn diagram of the identified upregulated DEGs for each time point. DEGs for the 1d time point are grouped in a green circle. DEGs for the 2d time point are grouped in a blue circle. DEGs for the 4d time point are grouped in a red circle. DEGs for the 7d time point are grouped in a yellow circle. DEGs for the 14d time point are grouped in an orange circle. Number of unique or shared DEGs are indicated.

Figure S7B: Venn diagram of the identified downregulated DEGs for each time point. DEGs for the 1d time point are grouped in a green circle. DEGs for the 2d time point are grouped in a blue circle. DEGs for the 4d time point are grouped in a red circle. DEGs for the 7d time point are grouped in a yellow circle. DEGs for the 14d time point are grouped in an orange circle. Number of unique or shared DEGs are indicated.

**Figure S8: Gene ontology analysis of enriched pathways for DEGs in the *R. norvegicus* samples.** Figure S8A: GO pathway analysis for upregulated DEGs shared for the 1d and 2d time points. Pathways were selected with a FDR of 0,05 and a minimum enrichment of 4. The name of each GO pathway is indicated on the left part of the figure. Bars represent the enrichment fold.

Figure S8B: GO pathway analysis for upregulated DEGs shared for the 7d and 14d time points. Pathways were selected with a FDR of 0,05 and a minimum enrichment of 4. The name of each GO pathway is indicated on the left part of the figure. Bars represent the enrichment fold.

Figure S8C: Western-blot showing the expression of Grm1 (VGLUT1) in infected or uninfected brain cell culture after 7 days. GAPDH is used as a loading control.

**Table S1:** **Number of DEGs identified by DESEQ2 for the *T. gondii* genome.**

**Table S2: Number of DEGs identified by DESEQ2 for the *R. norvegicus* genome.**

**Table S3:** **Gene expression for transcripts encoding proteins known to be involved in invasion.** Log_2_ fold change (FC) for each transcript at each time point of the infected brain cell culture. Color gradient depends on the value of FC. Downregulated values are represented shades of green. Upregulated values are represented in shade of red. Transcripts that were not detected are indicated by a double dash line (--). For each transcript the gene identification number (gene ID) and the corresponding annotation is also presented.
